# PEPATAC: An optimized pipeline for ATAC-seq data analysis with serial alignments

**DOI:** 10.1101/2020.10.21.347054

**Authors:** Jason P. Smith, M. Ryan Corces, Jin Xu, Vincent P. Reuter, Howard Y. Chang, Nathan C. Sheffield

## Abstract

**Motivation:** As chromatin accessibility data from ATAC-seq experiments continues to expand, there is continuing need for standardized analysis pipelines. Here, we present PEPATAC, an ATAC-seq pipeline that is easily applied to ATAC-seq projects of any size, from one-off experiments to large-scale sequencing projects.

**Results:** PEPATAC leverages unique features of ATAC-seq data to optimize for speed and accuracy, and it provides several unique analytical approaches. Output includes convenient quality control plots, summary statistics, and a variety of generally useful data formats to set the groundwork for subsequent project-specific data analysis. Downstream analysis is simplified by a standard definition format, modularity of components, and metadata APIs in R and Python. It is restartable, fault-tolerant, and can be run on local hardware, using any cluster resource manager, or in provided Linux containers. We also demonstrate the advantage of aligning to the mitochondrial genome serially, which improves the accuracy of alignment statistics and quality control metrics. PEPATAC is a robust and portable first step for any ATAC-seq project.

**Availability:** BSD2-licensed code and documentation at https://pepatac.databio.org.

## Introduction

Because cells package chromatin differently depending on their function and phenotype, profiling chromatin accessibility is a primary experimental approach for understanding cell states (1–3). The number of chromatin accessibility experiments has grown dramatically in recent years with the introduction of the assay for transposase-accessible chromatin (ATAC-seq) (4). With ATAC-seq now widespread, there is demand for analytical approaches (5, 6), including systematic processing pipelines to facilitate the goal of reproducible research and ease cross-study comparisons (7, 8).

To address this need we developed PEPATAC, a fast and effective ATAC-seq pipeline that easily generalizes across compute contexts and research environments. This pipeline has been built over years of experience analyzing chromatin accessibility experiments and implements several concepts that make it effective. These include ATAC-specific quality control outputs, both nucleotide-resolution and smoothed signal tracks, and a serial alignment strategy to deal with high mitochondrial contamination. Our serial alignment strategy, or ‘prealignments’, allows the user to configure a series of genomes to align to before the primary genome. PEPATAC provides a framework that allows a user to align serially in customized order to as many genomes as desired, which will be useful for many situations, including species contamination, dual-species experiments, repeat model alignments, decoy contamination, or spike-in controls.

While numerous ATAC-seq pipelines exist (For more in-depth coverage see: 5, 6), PEPATAC is designed with modularity and flexibility as paramount design considerations (Fig. 1a). PEPATAC is compatible with the Portable Encapsulated Projects (PEP) format (9), which defines a common project metadata description, allowing projects that use PEPATAC to be easily analyzed using any PEP-compatible tool. It also provides the possibility for a single project description to be shared across pipelines, computing environments, and analytical teams. PEPATAC is easily customizable, including changing individual command settings or even swapping specific software components by modifying a few lines of human readable configuration files.

**Fig. 1:**
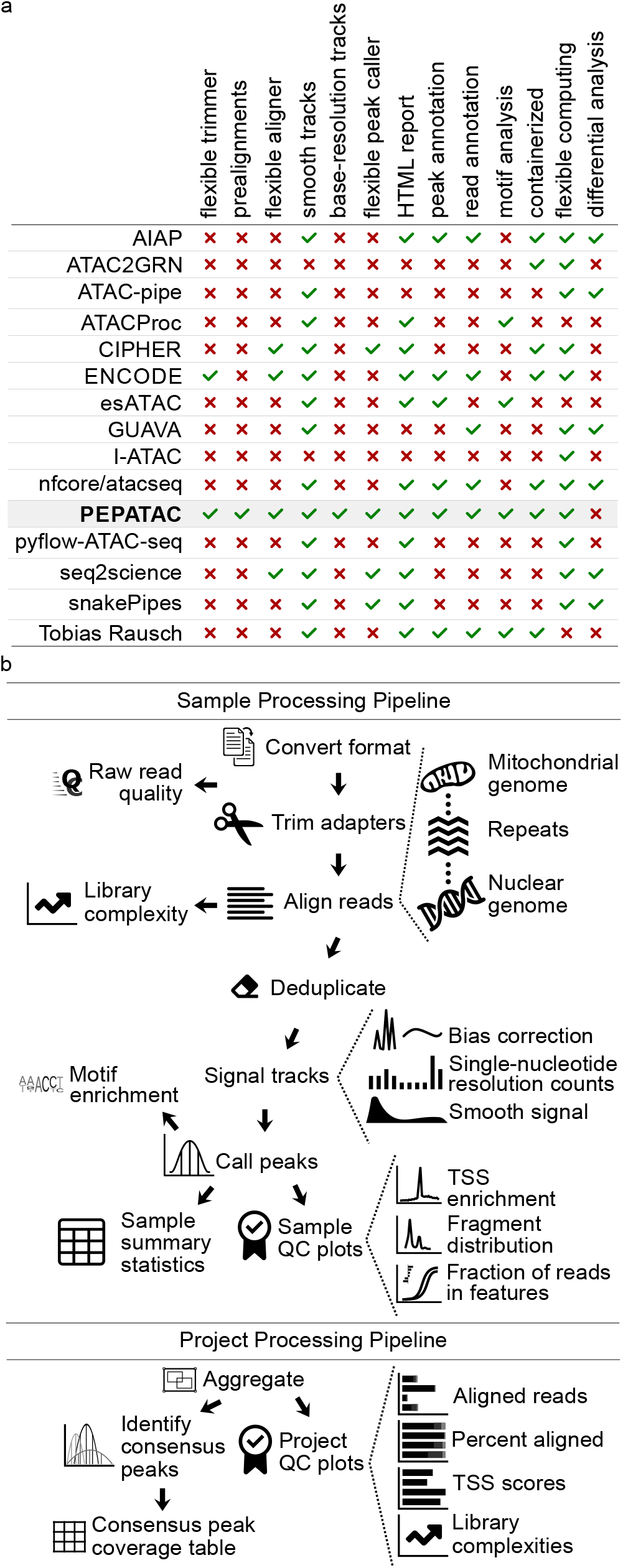
PEPATAC is feature-rich with a logical workflow. (a) We compared features across 14 ATAC-seq pipelines (AIAP (17); ATAC2GRN (18); ATAC-pipe (19); ATACProc (20); CIPHER (21); ENCODE (22); esATAC (23); GUAVA (24); I-ATAC (25); nfcore/atacseq (26); pyflow-ATAC-seq (27); seq2science (28); snakePipes (29); Tobias Rausch (30)) and PEPATAC stands out for being feature-rich. (b) Reads are preprocessed, serially aligned to the mitochondrial genome, curated repeats, and then the nuclear genome. PEPATAC generates both smooth and exact signal plots, called peaks, and QC output plots and tables.

PEPATAC does not rely on any specific local or cloud computing infrastructure, and it has already been deployed successfully in various compute environments at multiple research institutes to yield numerous peer-reviewed studies (10–14). While *all* ATAC-seq pipelines use several common bioinformatic tools (Fig. S1), we simplify the creation of a computing environment with the required command-line tools using conda (15), or either docker or singularity with the bulker multi-container environment manager (16).

PEPATAC includes a well-documented code base with detailed installation instructions, tutorials, and example projects, so it is useful for both the bench biologist and bioinformatician alike. We anticipate that this pipeline will provide a useful complete analysis for basic ATACseq projects and serve as a unified starting point for more advanced ATAC-seq projects.

## Materials and Methods

### PEPATAC configuration

The PEPATAC pipeline is divided into two major parts (Fig. 1b): First, it processes each sample individually at the *sample level.* Once sample processing is complete, the *project-level* part aggregates, analyzes, and summarizes the results across samples. PEPATAC is composed of two primary Python scripts that may be run from the command-line. Sample information and parameters are passed to the pipeline as command-line arguments (see `pepatac.py --help`), making it simple to use as a standalone pipeline for individual samples without requiring a complete project configuration. Project level output is produced using the project level pipeline (see `pepatac_collator.py --help`). PEPATAC is built using the Python module `pypiper` (31), which provides restartability, file integrity protection, copious logging, resource monitoring, and other features. Individual pipeline settings can also be configured using a pipeline configuration file (`pepatac.yaml`), which enables a user to specify absolute or relative paths to installed software, change adapter input files for trimming, and parameterize alignment and peak calling software tools. This configuration file comes with sensible defaults and will work out-of-the-box for research environments that include required software in the shell PATH, but it also may be configured to fit any computing environment and adapt to project-specific parameterization needs.

### Refgenie reference assembly resources

Like any genome analysis, PEPATAC relies on reference genome annotations. To ensure that results are comparable across runs, it’s important to use the same reference assembly. To manage these assets in a reproducible and robust manner, PEPATAC uses *refgenie*. `Refgenie` is a reference genome assembly asset manager that simplifies access to pre-indexed genomes and annotations for common assemblies, and also allows generating new standard reference genomes or annotations as needed while maintaining asset provenance (33, 34). For a complete analysis, PEPATAC requires several refgenie-managed assets: fasta, chrom_sizes, bowtie2_index, blacklist, refgene_tss, and feat_annotation. These can be either downloaded automatically or built manually, which require a genome fasta file, a gene set annotation file from RefGene, and an Ensembl gene and regulatory build annotation file. Using PEPATAC with `seqOutBias` requires the additional refgenie tallymemindex asset built for the same read length as the data. Many of these assets may also be directly specified at the command line should a user not have refgenie-managed versions available. The TSS annotation file, region blacklist, and feature annotation file may all be specified to use a local, user-specified file. For example, while ENCODE provides a common set of regions that are aberrantly overrepresented in sequencing experiments (e.g. a blacklisted set of regions) (35), a user may create their own version of regions that should be excluded from consideration and point to this file manually.

### File inputs and adapter trimming

PEPATAC sequentially trims, aligns, and analyzes sequences (Fig. 1b). PEPATAC accepts sequence data input in 3 formats: unaligned BAM, separated FASTQ, or interleaved FASTQ format. The pipeline first converts the input format into FASTQ (if necessary) for adapter trimming. For adapter trimming, users may select between skewer (36), trimmomatic (37), or an included Python tool using command-line arguments or the PEP configuration file. The pipeline stores quality control results including the number of raw, trimmed, or duplicated reads, and runs `FastQC` (38) if installed.

### Prealignments and mitochondrial DNA

Because ATAC-seq data can have a high proportion of reads mapping to the mitochondrial genome (from 15%-50% in a typical experiment up to 95% in some experiments (39)), we considered how to optimize the pipeline to deal with abundant mitochondrial DNA (mtDNA). High mtDNA exacerbates the alignment challenge caused by nuclear-mitochondrial DNA (NuMts), which are mtDNA sequences that have integrated into the nuclear genome throughout eukaryotic evolution (41). NuMts represent nonfunctional, truncated, and mutation-ridden copies of mitochondrial protein-coding genes; therefore, we assume that ATAC reads mapping to them are highly likely to be erroneous alignments. The typical strategy is to align to the mitochondrial and nuclear genomes simultaneously, and then remove nuclear-mitochondrial DNA (NuMts) post-hoc using a blacklist, but this suffers from three disadvantages: First, it is inefficient to align lots of mtDNA to the larger nuclear genome; second, reads that match both NuMt and mtDNA will be (incorrectly) split between the two, and third, this approach relies on an accurate pre-constructed annotation of NuMt locations, which may not be available for every reference genome. Furthermore, due to mitochondrial genetic diversity within and across cells, some reads derived from true mtDNA may in fact map better to the reference NuMt than to the reference mtDNA sequence. Also, reads that span the artificial breakpoint in the linear mtDNA reference may find an adequate NuMt match, but would never align to the mtDNA.

We found that by separately aligning first to the mitochondrial genome, we alleviated the challenges with simultaneous alignments. To capture NuMts that span the artificial breakpoint induced by converting the circular mitochondrial DNA into a linear representation for alignment, we use a doubled mitochondrial reference sequence, which enables non-circular aligners to align reads that span the breakpoint. By default, the pipeline is configured to align reads first to the doubled mitochondrial reference genome, but may be easily configured to perform any number of additional serial alignments.

### Alignments, deduplication, and library complexity

For prealignments and primary alignment, PEPATAC employs bowtie2 by default (42). Bowtie2 settings are configurable in the pipeline configuration file but come with sensible defaults of `-k 1 -D 20 -R 3 -N 1 -L 20 -i S,1,0.50` for prealignments and `--very-sensitive -X 2000` for nuclear genome alignment. Users may optionally use bwa (43) with settings similarly configurable in the pipeline configuration file (default: `-M`). Following alignment, reads with mapping quality scores below 10 and any residual mitochondrial reads are removed and read deduplication is carried out using samblaster (44), but picard’s MarkDuplicates (45), or samtools (46) may also be utilized based on user preference. PEPATAC utilizes *preseq* (47) to calculate and plot sample library complexity at the current depth, and includes the number of independently calculated duplicates (Fig 2a). The pipeline also projects the unique fraction of the library at 10M total reads. These metrics provide an estimate of library complexity and allow the user to determine the value of subsequent sequencing.

**Fig. 2:**
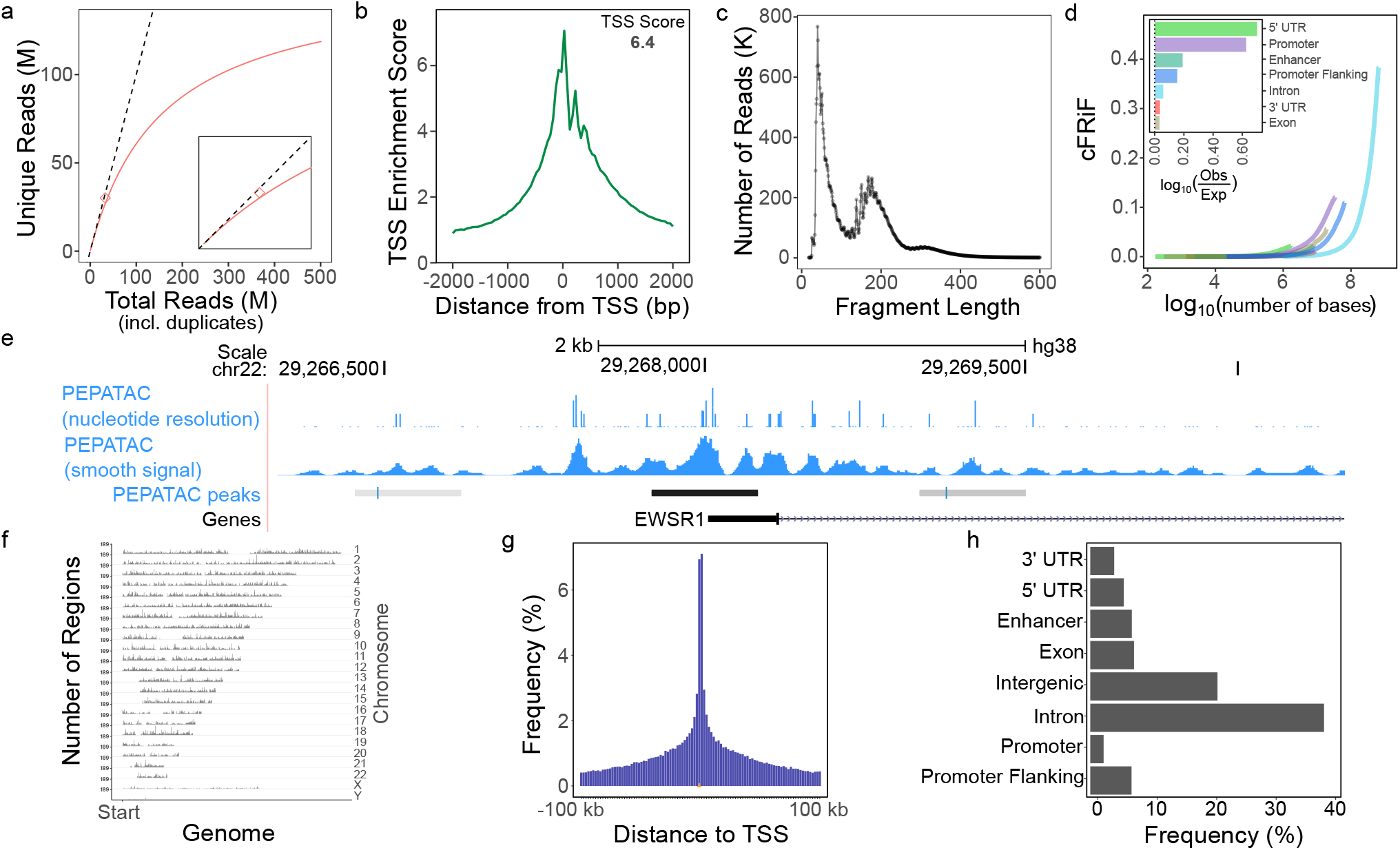
Example PEPATAC QC plots for reads and peaks. (a) Library complexity plots the read count versus externally calculated deduplicated read counts. Red line is library complexity curve for SRR5427743. Dashed line represents a completely unique library. Red diamond is the externally calculated duplicate read count. (b) TSS enrichment quality control plot. (c) Fragment length distribution showing characteristic peaks at mono-, di-, and tri-nucleosomes. (d) Cumulative fraction of reads in annotated genomic features (cFRiF). Inset: Fraction of reads in those features (FRiF). e) Signal tracks including: nucleotide-resolution and smoothed signal tracks. PEPATAC default peaks are called using the default pipeline settings for MACS2 (32). (f) Distribution of peaks over the genome. (g) Distribution of peaks relative to TSS. (h) Distribution of peaks in annotated genomic partitions. Data from SRR5427743.

### Library QC metrics

For quality control, PEPATAC provides a TSS enrichment plot, produced by aggregating reads present in regions 2000 bases upstream and downstream of a reference set of TSSs (Fig 2b). Enrichment is calculated as the average number of reads in a 100 bp window around the TSS divided by the average number of reads in the first 200 bases of the entire region. This yields low signals in the tails with a peak in the center, which we take to be the TSS enrichment score. PEPATAC also produces a fragment length distribution plot (Figure 2c). A standard quality ATAC-seq library is expected to yield clearly defined peaks at open chromatin (<100bp), mononucleosomes (200 bp), and sequentially smaller peaks representing multi-nucleosomes at regular intervals. To evaluate the enrichment of all reads across genomic partitions, PEPATAC plots both the fraction and cumulative fraction of reads (FRiF, cFRiF respectively) in genomic features (Fig 2d). A novel feature of PEPATAC includes the plotting of the fraction of reads in any feature type, not solely in peaks. This is plotted as the cumulative sum of reads in each feature divided by the total number of aligned reads against the cumulative sum of bases in each feature. The relative proportion of each feature can be then be directly compared. The standard feature annotation produced and managed by `refgenie` includes Ensembl defined enhancers, promoters, promoter flanking regions, 5’ UTR, 3’ UTR, exons, and introns in that order. Users can specify an alternative annotation file, either a custom one or simply a different sort order, using the `--anno-name` pipeline parameter. For a quality sample, the proportion of reads in peaks should be the most enriched, reflecting the specificity of the peak calls for that sample.

### Signal tracks and peak calling

Alignments are used to generate two signal tracks: one that records the exact location of transposition events, and one that is smoothed (Fig 2e). These tracks may be used for different downstream analyses; the exact track is useful for analysis that requires nucleotide-resolution, while the smoothed version is often preferred for visualization and peak analysis. Reads, representing transpoase cut-sites, are extracted from the deduplicated, low-quality removed, primary genome mapped BAM file into a wiggle-like track. For the exact signal track, these cut-sites are shifted +4 bases for positive strand reads and −5 bases for negative strand reads. For the smooth signal track, we extend the shifted exact sites +/− 25 bases to yield 50 bp smoothed windows around the exact cut-site position. seqOutBias is an optional tool that can be used to correct for enzymatic (e.g. Tn5 transposase) bias and generate tracks for visualization (48). The bias itself is corrected using a k-mer mask for the plus and minus strand Tn5 recognition sites and by taking the ratio of genome-wide observed read counts to the expected sequence based counts for each k-mer (48). The k-mer counts take into account mappability at a given read length using GenomeTools’ Tallymer program (49).

An earlier study found multiple peak callers worked well with chromatin accessibility data (50), and PEPATAC provides the option to use F-Seq (51), MACS2 (32), Genrich (52), HOMER (53), or HMMRATAC (54) for peak calling, with parameters customizable in the pipeline configuration file. MACS2 is used by default (`--shift −75 --extsize 150 --nomodel --call-summits --nolambda --keep-dup all -p 0.01`). The default settings are intended to maximize recall and sensitivity. More stringent settings can be easily adopted by modifying the pipeline configuration file. Called peaks are standardized by extending up and down 250 bases (a tunable parameter, `--extend`) from the summit of each peak to establish peaks 500 bases in width. Any peaks which then extend beyond chromosome boundaries are trimmed. Utilizing fixed-width peaks reduces bias towards larger peaks in both count-based and motif analyses while simultaneously improving the identification of consensus peak sets by reducing the likelihood of extraordinary large peaks created through the union and merging of multiple peak sets. Finally, peak scores are normalized to score per million by dividing by the sum of scores over 1M.

PEPATAC also produces several plots detailing enrichment of reads in peaks including: the distribution of peaks across the genome by chromosomal location (Fig 2f), the distribution of peaks relative to TSSs (Fig 2g), and the distribution of peaks within genomic partitions (Fig 2h). The TSS distance distribution shows the distance of called peaks with respect to TSSs grouped in log-scale bins. Finally, users may optionally employ HOMER to calculate motif enrichments in called peaks (55).

### Running multiple samples with PEPATAC

To run the pipeline across multiple samples in a larger project, the pipeline uses the job submission engine `looper` (56), which employs the Portable Encapsulated Project standardized definition of project metadata (9)(Fig. S2). This standard project format enables a pipeline to be run on any project that follows the format, which is simple, standardized, and well-documented. `Looper` enables the PEPATAC pipeline to be run in any compute environment, including locally (the default) on a single laptop or desktop, or with any cluster resource manager. It also can be used with containers. Addi-tionally, `looper’s` project format gives pipeline users access to APIs written in Python and R for downstream analysis of pipeline results.

For the user whose environment is set up to run containers, we enable container use with either Docker or Singularity via a single image file or through the multicontainer environment manager, `bulker` (16). Using `bulker`, PEPATAC may be run in containers across samples and compute environments, simplifying deployment by requiring only `bulker` and the PEPATAC pipeline itself, eliminating the need to install each required package independently.

### Aggregating results from multiple samples

To summarize and incorporate data across samples, the second step in a PEPATAC analysis is to run a projectlevel pipeline (`pepatac_collator.py`) that identifies consensus peaks across a project and calculates sample coverage of those consensus peaks in a convenient table for easy downstream analysis. To establish consensus peaks, PEPATAC identifies overlapping (1 bp, a tunable parameter: `--min-olap`) peaks between every sample in a project and defines the consensus peak’s coordinates based on the overlapping peak with the highest score. Peaks present in at least 2 (parameter: `--cutoff`) samples with a minimum score per million greater than or equal to 5 (parameter: `--min-score`) are retained. A peak count table is then provided where every sample peak set is overlapped against the consensus peak set. Individual peak counts for an overlapping peak are weighted by multiplying by the percent overlap of the sample peak with the consensus peak.

For navigating results, PEPATAC provides both sample and project level reports in a convenient, easy-to-navigate HTML report with project-level summary table and plots, job status page, and individual sample pages with sample statistics and QC plots all at your fingertips. In addition, `looper` will produce summary plots from individual sample statistics including the number of aligned reads, percent aligned reads, TSS scores, and library complexities. A user can produce the HTML report during a run or after completion, with the job status page providing information on whether a sample has failed, is still running, or has already completed.

## Results

To demonstrate PEPATAC’s default workflow and output, we analyzed samples from the original standard ATAC (4), fast ATAC (57), and omni ATAC (58) protocol papers. This dataset includes human ATAC-seq reads from 33 standard ATAC, 152 fast ATAC, and 139 omni ATAC samples (Supplemental file 1). PEPATAC provides output and quality control results both for individual samples and for the project as a whole. For each sample, PEPATAC produces narrowPeak and bigWig files to visualize nucleotide-resolution alignments, smoothed alignments, and peak calls. PEPATAC also produces summary statistics files that report the number of reads, duplicates, genome alignment rates, transcription start site (TSS) enrichment score, number of called peaks, fraction of reads in peaks (FRiP), and job runtime among others for every sample in a project.

### Performance

PEPATAC is designed to be computationally efficient. To evaluate how PEPATAC scales with increasing numbers of reads, we ran 430 ATAC-seq samples of varying input size through PEPATAC (Supplemental file 4). We then placed samples in 500MB input file size bins and compared runtimes and peak memory usage (Fig. S3). Runtime scales linearly with increasing file size, but importantly, even samples with more than 150 million reads completed in less than 8 hours (Fig S3a). We also show that PEPATAC, with default settings, only utilizes between 5-9 GB at peak memory use (Fig S3b).

### Prealignments

To evaluate the advantage of serially aligning to the mitochondrial genome (Fig. 3a), we measured the total alignment runtime of synthetic mixtures of mitochondrial-aligning (mtDNA) and whole human-aligning (hg38) sequences with and without prealignments. We constructed libraries of mixed mtDNA:hg38 mapping ATAC-seq reads from 0% to 100% mtDNA in increments of 10%, at 10 million, 20 million, and up to 200 million total reads in increments of 20 million reads, resulting in 121 different library combinations. We recorded the alignment time for each input file with and without prealignments (Fig. 3b). To determine for which scenarios using prealignments is beneficial, we calculated the log ratio of run times with prealignments versus without prealignments and found that using prealignments reduces the total time of alignment even when mtDNA alignment rates are under 10% (Fig. 3c). In addition to speed and efficiency gains, PEPATAC with prealignment compared to without prealignment to mtDNA yields higher alignment rates to mitochondrial sequence than aligning to a combined human and mitochondrial genome as is commonly performed (Fig. 3d). This is true for every sample tested no matter the library preparation protocol nor percent mitochondrial contamination (Fig. S4). This result indicates that the common approach of simultaneously aligning to the nuclear and mitochondrial genomes systematically underestimates the fraction of mitochondrial reads in an experiment. We therefore propose that mitochondrial alignment rates are generally underestimated by about 1-5% in published reports.

**Fig. 3:**
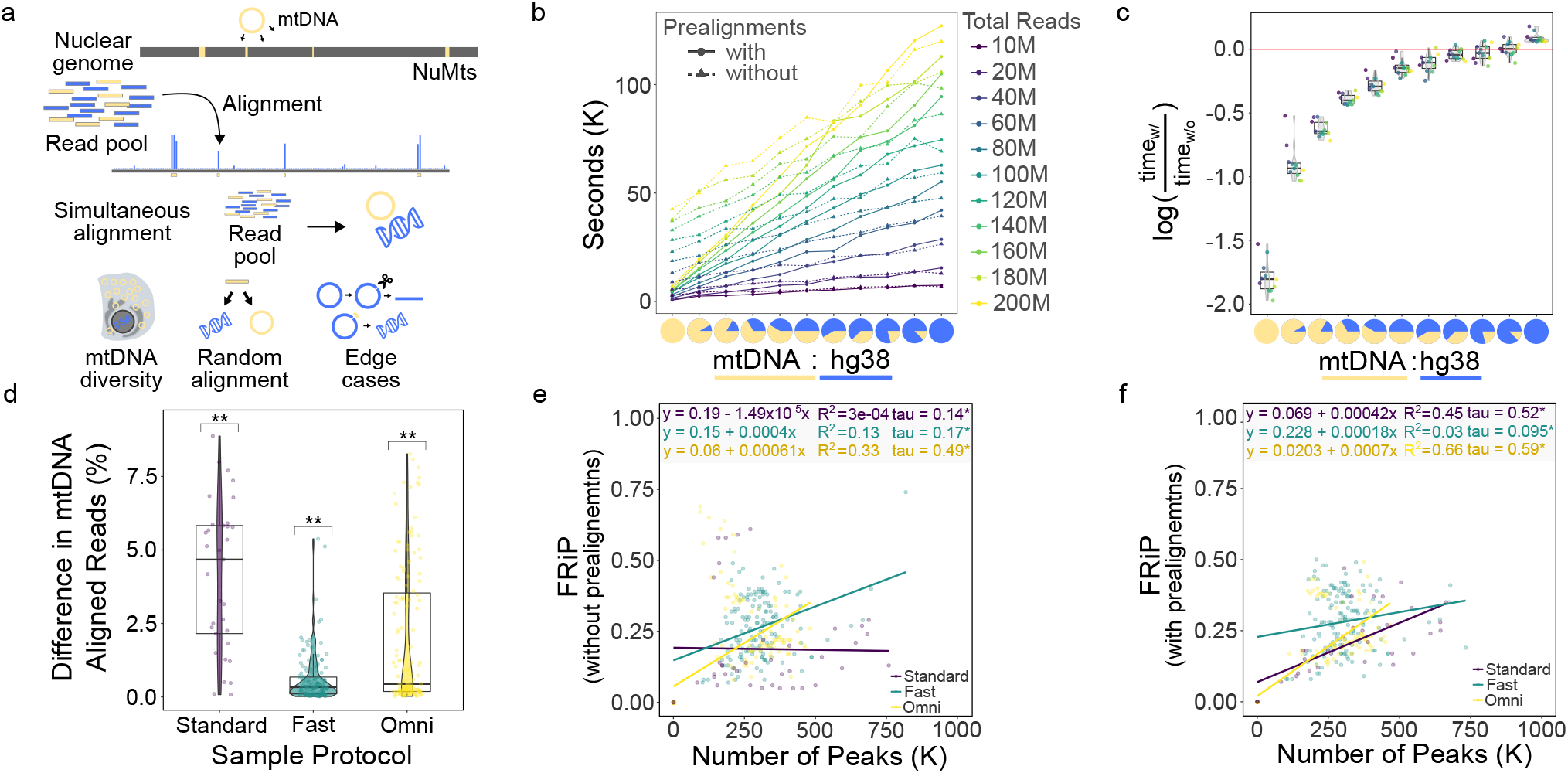
PEPATAC prealignments increase mapped mtDNA reads, improve computational efficiency, and positively influences the fraction of reads in peaks (FRiP) metric. (a) NuMTs represent a significant complication of simultaneous alignment. (b) At mtDNA percentages from 10-100% at total read numbers ranging from 10-200M, using prealignments dramatically reduces run time. (c) Log ratio of prealignments runtimes versus no prealignment runtimes yields significant savings. (d) There is a significant increase in the percent of reads mapped to mitochondrial sequence when using prealignments versus not across standard, fast, and omni-ATAC protocols. (e) As reported for ChIP-seq (59), FRiP is positively correlated with the number of called peaks. (f) With prealignments, the positive correlation between FRiP and the number of called peaks tends to increase ((d) ** = p < 0.001; t-test (mu = 0) with Benjamini-Hochberg correction. (e-f):* = p < 0.0001; Kendall rank correlation coefficient).

To show how prealignments successfully depletes reads aligning to NuMTs, we ran a standard ATAC (SRR5427804), fast ATAC (SRR2920492), and omni ATAC (SRR5427806) sample through PEPATAC with no prealignments, prealignment to mitochondrial sequence, and prealignment to mitochondrial, ribosomal, and known repeat sequences. We then compared the highest signal peaks between each prealignment strategy across each ATAC-seq protocol. We used BLAST (60) to annotate the highest signal peaks and then intersected called peaks under each strategy with the ENCODE blacklist (35), which normally is used to filter results in PEPATAC by default. The omni ATAC sample had the least number of aberrant high signal peaks with only a single NuMT peak identified in the top 10 highest signal peaks and *only* present when analyzed without prealignments. Significantly, as soon as mitochondrial prealignment is included, this peak is excluded (Supplemental file 3, Fig. S5a). Of the top 100 omni ATAC peaks, there are fewer overlaps with blacklisted regions, both overall, and as we increase the number of prealignments. With no prealignments there are 4 blacklisted regions in the top 100 and only 2 with prealignments (Supplemental file 3). As omni ATAC is reported to reduce mitochondrial reads, this result is expected. Furthermore, this difference is highlighted as we compare both fast ATAC and standard ATAC. Three of the top 10 peaks from the fast ATAC sample without prealignments aligned to mitochondrial sequence (Supplemental file 3). These are eliminated with prealignments. Additionally, without prealignments, 22 of the top 100 peaks intersect blacklisted regions. Only 18 overlap with mitochondrial prealignment, and significantly, only 3 of the top 100 overlap blacklisted regions when prealigning includes ribosomal and repeat regions (i.e. satellite DNA). This suggests that a number of regularly identified peaks should typically be excluded in the absence of prealignments. While a blacklist does an excellent job at removing these regions, prealignment achieves similar results while also removing additional non-blacklisted regions that are likely spurious (mapping to unmapped regions or to different species, see Supplemental file 3). These results are even more obvious with standard ATAC. Standard ATAC without prealignment to mitochondria mapped 8 of the top 10 peaks to NuMTs (Supplemental file 3). These are removed with prealignment to mitochondria. Furthermore, the number of blacklisted regions drops from 17 without prealignments to 7 with mitochondrial prealignment and only 2 with mitochondrial, ribosomal, and repeat region prealignment. Because prealignment reduces spurious peak assignment (Supplemental file 3, Fig. S5b) *and* it reduces total runtime in nearly every scenario (Fig. 3c), prealignment is an effective strategy to include in every pipeline run.

### Peak caller comparison

To evaluate the difference in called peaks when using different peak callers, we compared both the PEPATAC determined consensus peaks and the peaks from a single sample (SRR5210416) produced when using different peak callers (Fseq, Genrich, HOMER, HMMRATAC, MACS2 with variable peaks, and MACS2 with fixed peaks). Similarity between the intervals was evaluated with a modified Jaccard statistic (61) implemented in the bedtools (62) package. At the single sample level MACS2 with variable peak width is the most similar in output to MACS2 with fixed peaks and Fseq (Fig. S6a, see Supplemental file 2). Interestingly, the least similar peak results are from Genrich and HMMRATAC, which possibly reflects the goal of both tools being designed to evaluate ATAC-seq data as opposed to originally being developed for ChIP-seq (Fig. S6a). These differences become more pronounced at the consensus peak level, with HMMRATAC becoming more dissimilar (average jaccard statistic = 0.31, Supplemental file 2) to the other peak callers (Fig. S6b).

We also asked whether this difference was due to an improvement in reduced peak calling at nuclear mitochondrial sequences (NuMTs), repeat regions, or high signal regions. One way to evaluate this is to determine the number of intersections of the individual peak caller called regions against a known blacklist (35) and to BLAST (60) the highest signal peaks. Indeed, HMMRATAC overlaps the least number of blacklisted regions (231 versus the maximum of 756 with HOMER; see Supplemental file 2) and it turns out a number of both the blacklisted regions and the highest signal peaks are NuMTs or repeat regions (Supplemental file 3). While MACS2 remains the most commonly employed peak caller across ATAC-seq pipelines, further comparative studies may better illustrate the utility of some of the more recently developed peak callers.

### Library QC comparison

Several of the QC metrics (e.g. TSS enrichment score, the fragment distributions, non-redundant fractions, and the PCR bottlenecking coefficients 1 and 2) employed by PEPATAC are near-universal in the field, and as such are calculated in the same manner. To evaluate how different annotations may affect the TSS score, we also compared TSS annotations from Ensembl, Gencode, and Refgene (PEPATAC default). Refgene produces higher TSS scores (Fig. S7), which reflects the fact that Refgene contains only the most commonly employed transcription start sites for each gene whereas both Ensembl and Gencode include all known sites, diluting the aggregated signal.

### Fraction of reads in peaks

It has also been reported that in ChIP-seq experiments, but not specifically in ATAC-seq, that FRiP correlates positively with the number of identified peaks (59) (Fig. 3e). In libraries with significant mitochondrial contamination, for example, from libraries produced using standard-ATAC library preparation protocols, this correlation is emphasized when using prealignments (Fig. 3f). We next sought to understand how the serial alignment strategy affects calculation of Fraction of Reads in Peaks (FRiP). FRiP is a common qualitative measure of enrichment and sample quality. However, FRiP calculations are poorly defined, making it dangerous to compare FRiP scores among different protocols and approaches. ENCODE defines the denominator of the FRiP score to be total mapped reads (ENCODE Terms). If only one genome is used for alignment, then the calculation is clear, but for a serial alignment pipeline, the FRiP score depends on whether the denominator includes reads mapped to the nuclear genome only, or to all genomes (Fig. S8c,d). By default, PEPATAC uses the deduplicated, low-quality removed, primary genome mapped BAM file to calculate the fraction of reads in the final called peak output file, which by default utilizes fixed width peaks and has removed any blacklisted regions. This has the consequence of changing the FRiP calculation based on whether prealignments were used (Fig. S8c,d). When using prealignments, the default FRiP calculation will significantly increase, because the number of reads mapped to the primary genome is reduced due to reads mapping more accurately to the mitochondrial genome and thus being excluded from downstream analysis. When FRiP is calculated using the total mapped reads (prealignments **and** primary alignment), these relationships are inversed (Fig. S8c,d). In any scenario, prealignments lead to more total mapped reads, due to more efficient mitochondrial alignment. As more recent ATAC-seq sample preparation protocols intentionally reduce mitochondrial contamination, these differences are most pronounced when using the original, standard ATAC-seq protocol. Therefore, reliance on a specific cutoff (e.g. 0.3 or greater) as indicative of a quality sample must be relative to protocol and method.

## Discussion

PEPATAC is an efficient, user-friendly ATAC-seq pipeline that produces helpful quality control plots and signal tracks that provide a comprehensive starting point for further downstream analysis. Two key benefits of the PEPATAC pipeline over existing pipelines are its flexibility and modularity. PEPATAC is uniquely flexible, for example, by allowing pipeline users to serially align to multiple genomes, to select from multiple aligners, peak callers, and adapter trimmers, while providing a convenient, configurable interface so a user can adjust parameters for individual pipeline tasks. Furthermore, PEPATAC reads projects in PEP format, a standardized, well-described project definition format, providing a reproducible interface with Python and R APIs to simplify downstream analysis.

Because PEPATAC is built on `looper`, it is easily deployable on any compute infrastructure, including a laptop, a compute cluster, or the cloud. It is thereby inherently expandable from single to multi-sample analyses with both project level and individual sample level quality control reporting. This means that a user may submit any number of samples using *a single looper command and corresponding PEP metadata file.* Its design allows for simple restarts at any step in the process should the pipeline be interrupted. Due to its modular construction multiple software options for primary pipeline steps are available, creating a swappable pipeline flow path with individual steps adaptable to future changes in the field. PEPATAC is a rapid, flexible, and portable ATAC-seq project analysis pipeline providing a standardized foundation for more advanced inquiries.

## Documentation and links

- PEPATAC v0.9.16: pepatac.databio.org.
- PEP metadata standards: pep.databio.org.
- `Looper` job submission engine: looper.databio.org.
- `Refgenie` reference genomes: refgenie.databio.org.
- Source code to reproduce output for this paper: github.com/databio/pepatac_paper_data.

## Acknowledgments

We would like to acknowledge helpful discussions and contributions from Yuning Wei, Ying Shen, and Jason Buenrostro on early versions of the pipeline. This work was supported by NIH grants RM1-HG007735 (to H.Y.C.) and 1R35GM128636-01 (to N.C.S.). H.Y.C. is an Investigator of the Howard Hughes Medical Institute. M.R.C. is supported by a K99 (K99-AG059918) award and is an American Society of Hematology Scholar.

## Declaration

H. Y.C. is a co-founder of Accent Therapeutics, Boundless Bio, and a consultant for Arsenal Biosciences and Spring Discovery. Stanford University holds a patent on ATAC-seq on which H.Y.C. is a named inventor.

## Supplemental figures

**Fig. S1:**
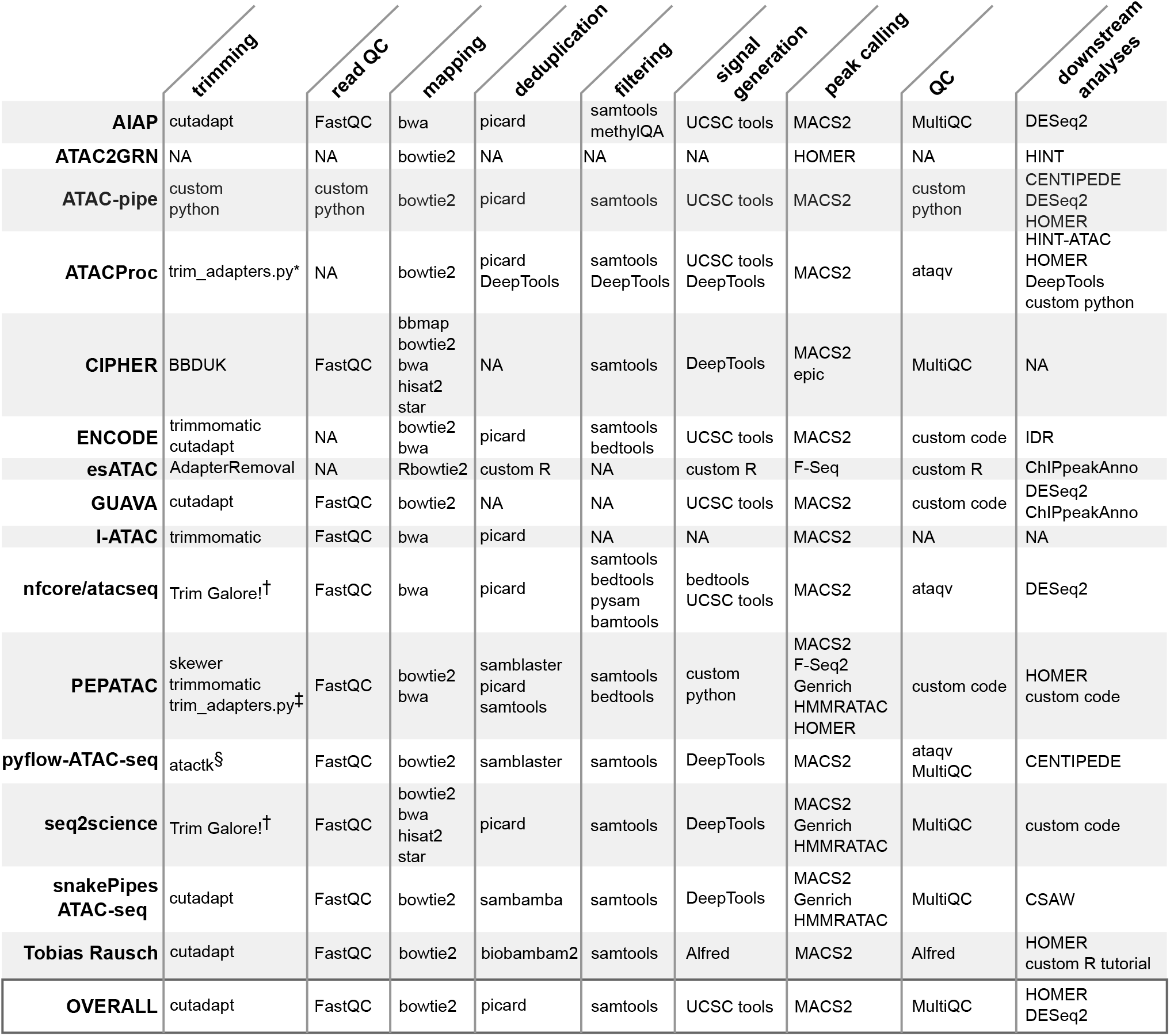
ATAC-seq pipelines universally require several common bioinformatic tools. While all pipelines require a number of common bioinformatic tools, PEPATAC offers the greatest flexibility and includes a number of the most popular tools.

## Supplemental files

### Supplementalfile1.csv

Supplemental_file_1.csv is the PEP-formatted sample table for the primary dataset. Samples are defined by protocol, whether standard, fast, or omni, and include accession numbers for access through the Gene Expression Omnibus (63).

### Supplementalfile2.xlsx

Supplemental_file_2.xlsx contains two sheets. The “jaccard_similarities” sheet includes tables representing the results of `bedtools intersect` between each independent peak caller software for 1) the PEPATAC derived consensus peak set, and 2) for an individual sample (SRR5210416) between each peak caller. This sheet also includes the average jaccard statistic for each peak caller. The “blacklisted_regions” sheet compares the number of peaks generated by each peak caller that overlap blacklisted regions (35).

### Supplementalfile3.xlsx

Supplemental_file_3.xlsx includes three sheets for a standard ATAC (SRR5427804), fast ATAC (SRR2920492), and omni ATAC (SRR5427806) sample that has been run through PEPATAC with 1) no prealignments, 2) mitochondrial prealignment (rCRSd: the revised Cambridge Reference Sequence doubled genome), and 3) mitochondrial, human repeats, and rDNA prealignments. In each sheet, for the highest scoring peaks, individual peak fasta sequences (included) were aligned with BLAST (60) and top scoring annotations recorded. If the peak overlaps a known blacklisted region (35), this is also marked.

**Fig. S2:**
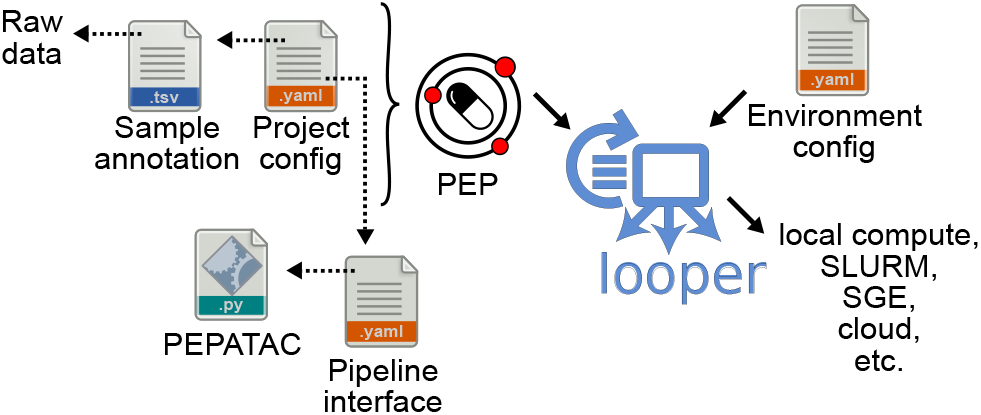
Deploying PEPATAC across multiple samples using looper. The PEPATAC pipeline can be easily run across multiple samples in any computing environment using looper.

**Fig. S3:**
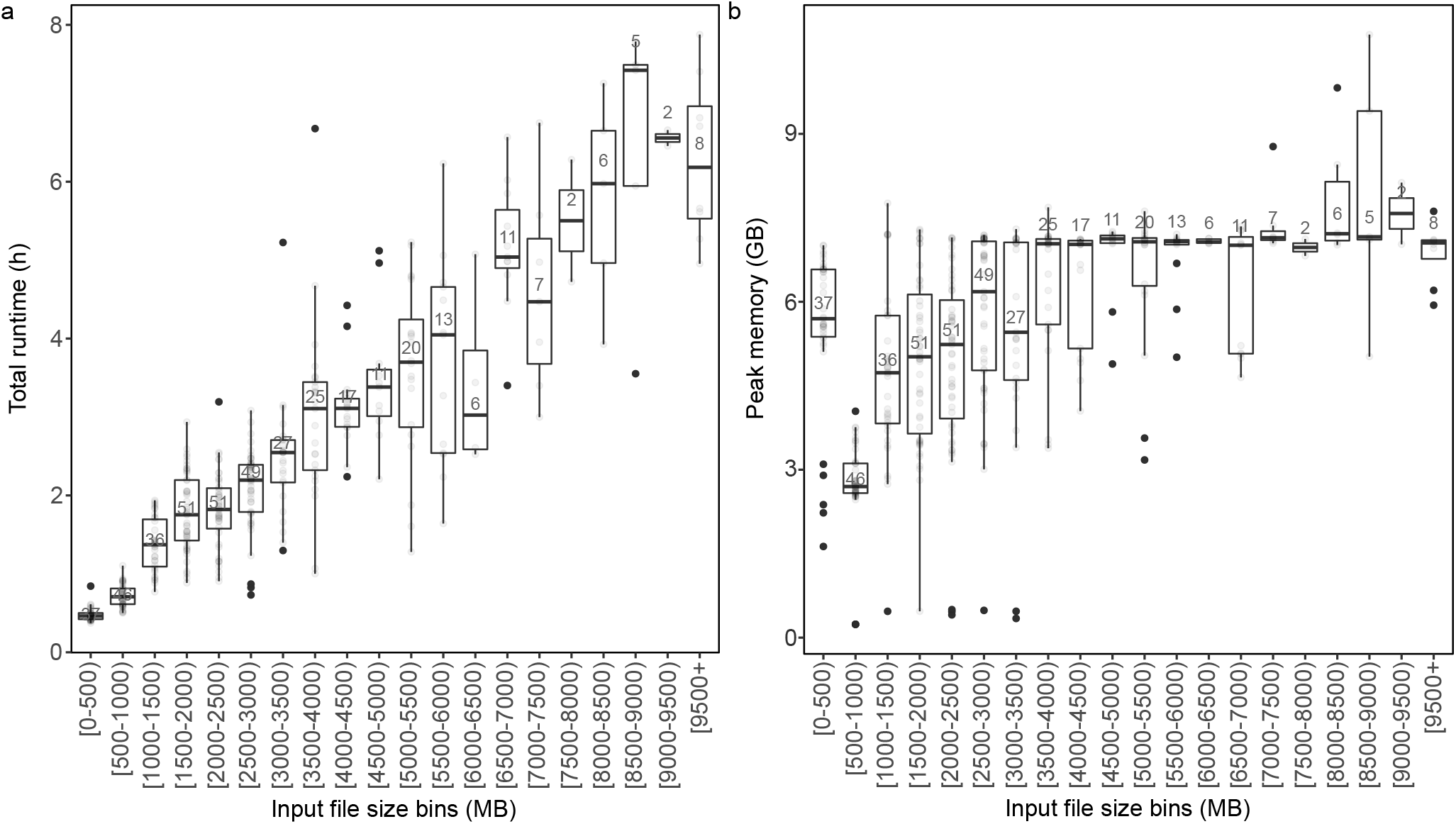
PEPATAC is computational efficient. (a) Pipeline runtime scales linearly with input file size. (b) Pipeline memory use peaks between 5-9GB.

### Supplementalfile4.csv

Supplemental_file_4.csv is the PEP-formatted sample table for the performance testing dataset. Accession numbers for file access through the Gene Expression Omnibus (63) are included for each sample.

**Fig. S4:**
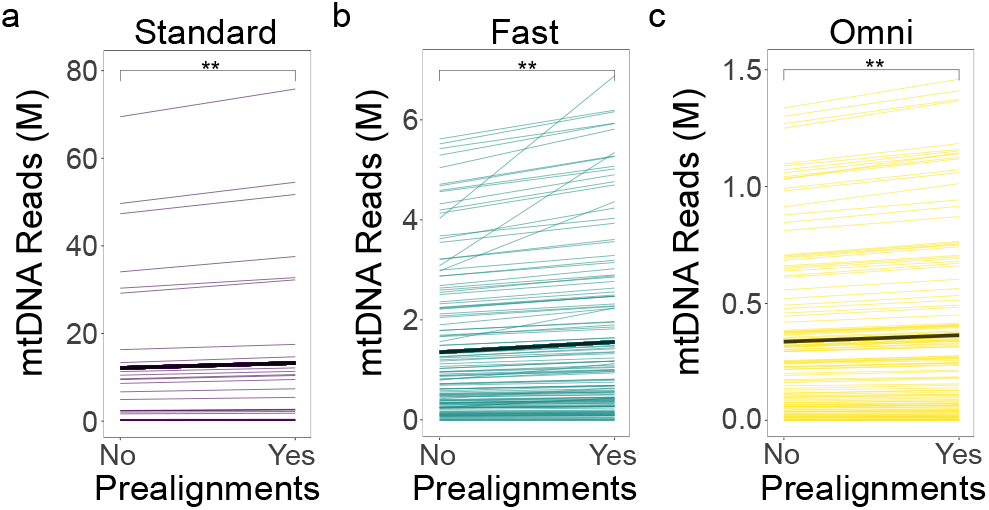
Prealignment increases mtDNA alignment. Within Standard (a), Fast (b), and Omni (c) ATAC-seq library preparation protocols, every sample shows increased mtDNA alignment when utilizing prealignments (The gray lines represent the mean increase within each protocol. ** = p < 0.001; t-test (mu = 0) with Benjamini-Hochberg correction.)

**Fig. S5:**
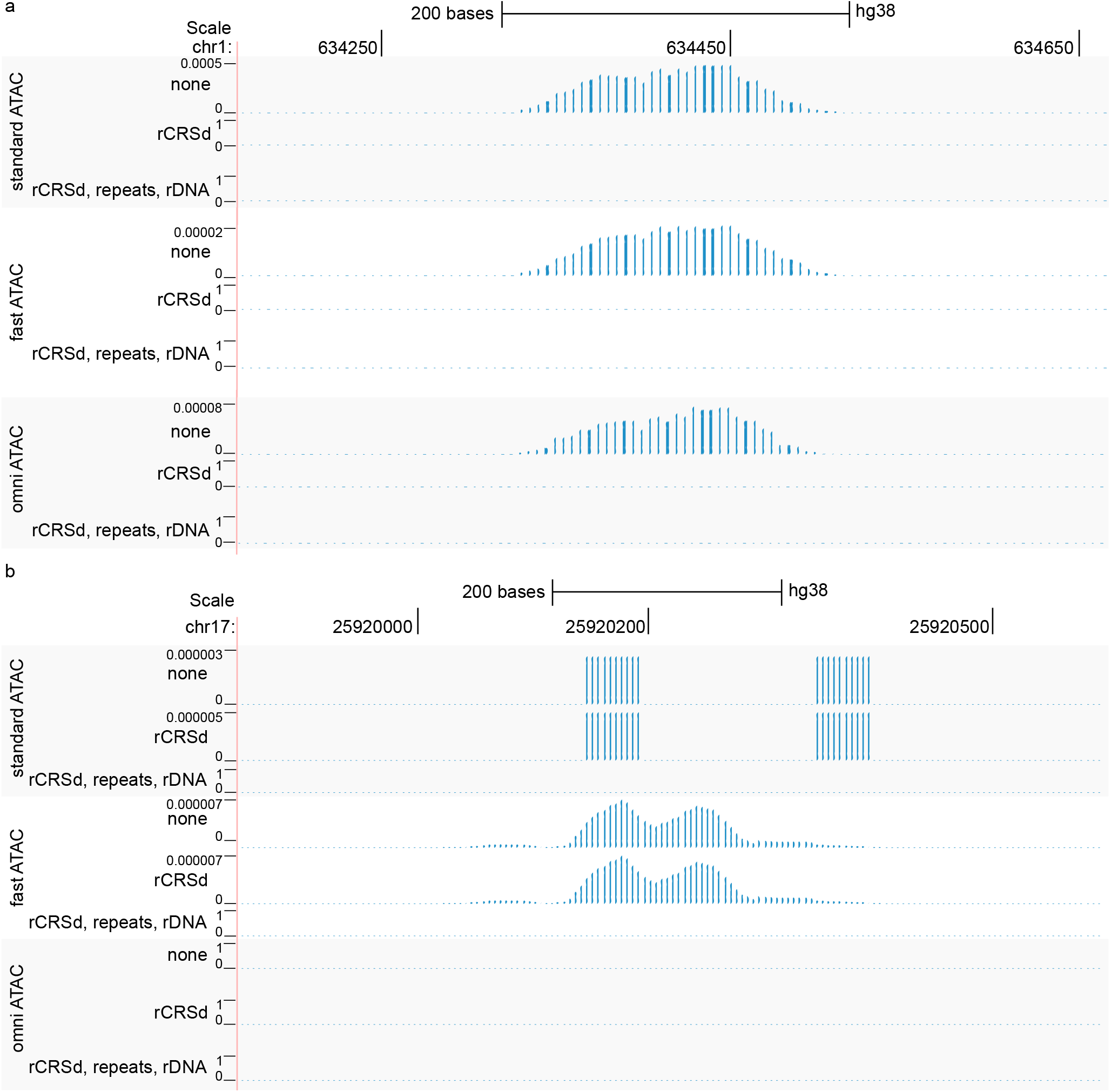
Prealignment (and improved ATAC-seq library preparation protocols) successfully deplete signal from NuMTs, repeat regions, and high signal regions. (a) Even where improved library preparation protocol leads to a NuMT annotated peak, prealignment successfully removes the spurious signal. (b) Both omni ATAC and prealignment to mitochondria and repeats and ribosomal sequence successfully depletes a spurious signal.

**Fig. S6:**
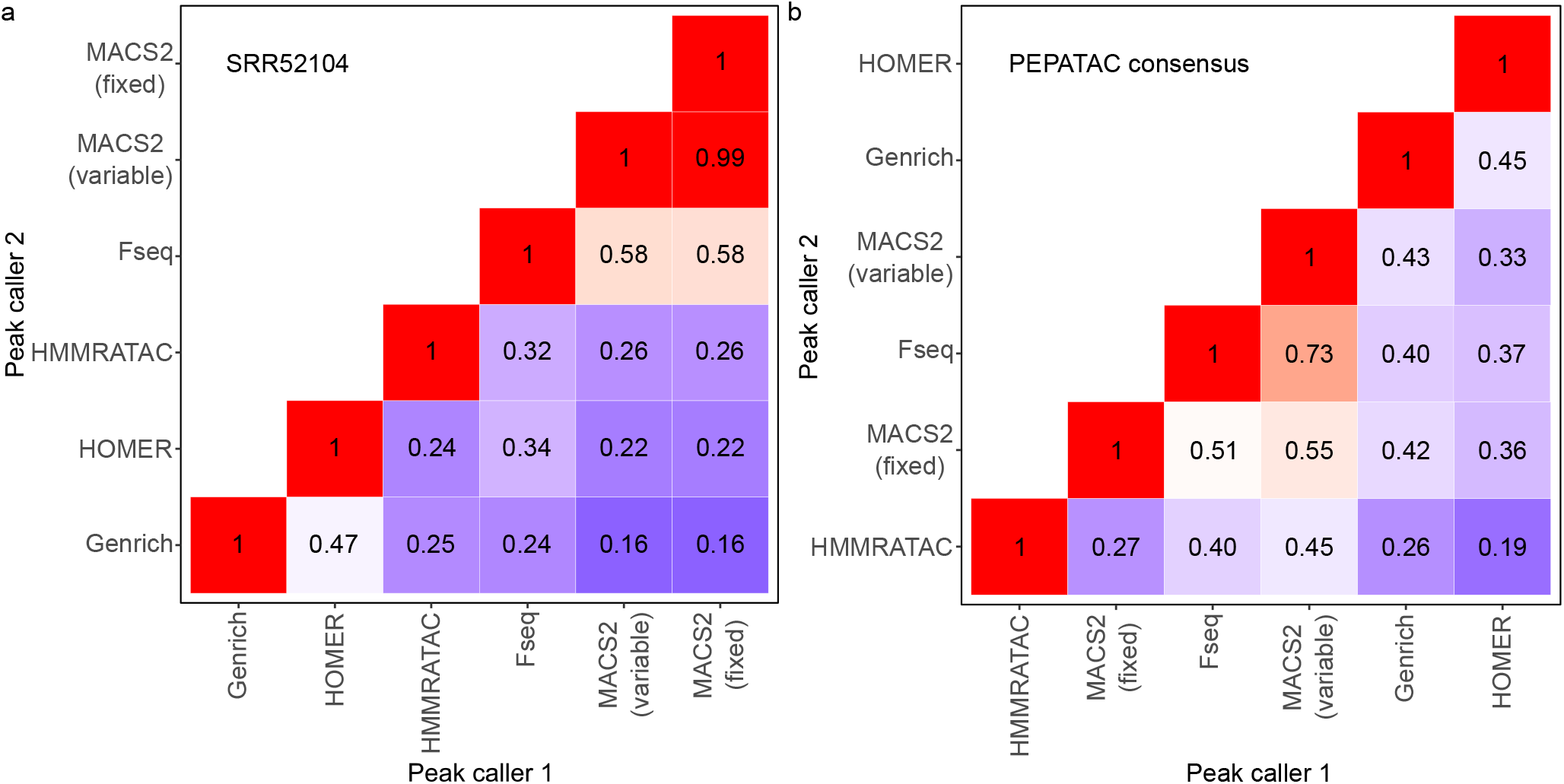
Peaks are comparatively dissimilar between the five optional peak callers. (a) For a single sample, MACS2 derived peaks, both with fixed and variable width peaks, are the most similar to Fseq called peaks. Genrich and HMMRATAC are the most unique among peak callers. (b) After PEPATAC consensus peak generation, HMMRATAC becomes even more dissimilar from the results derived from alternative peak callers.

**Fig. S7:**
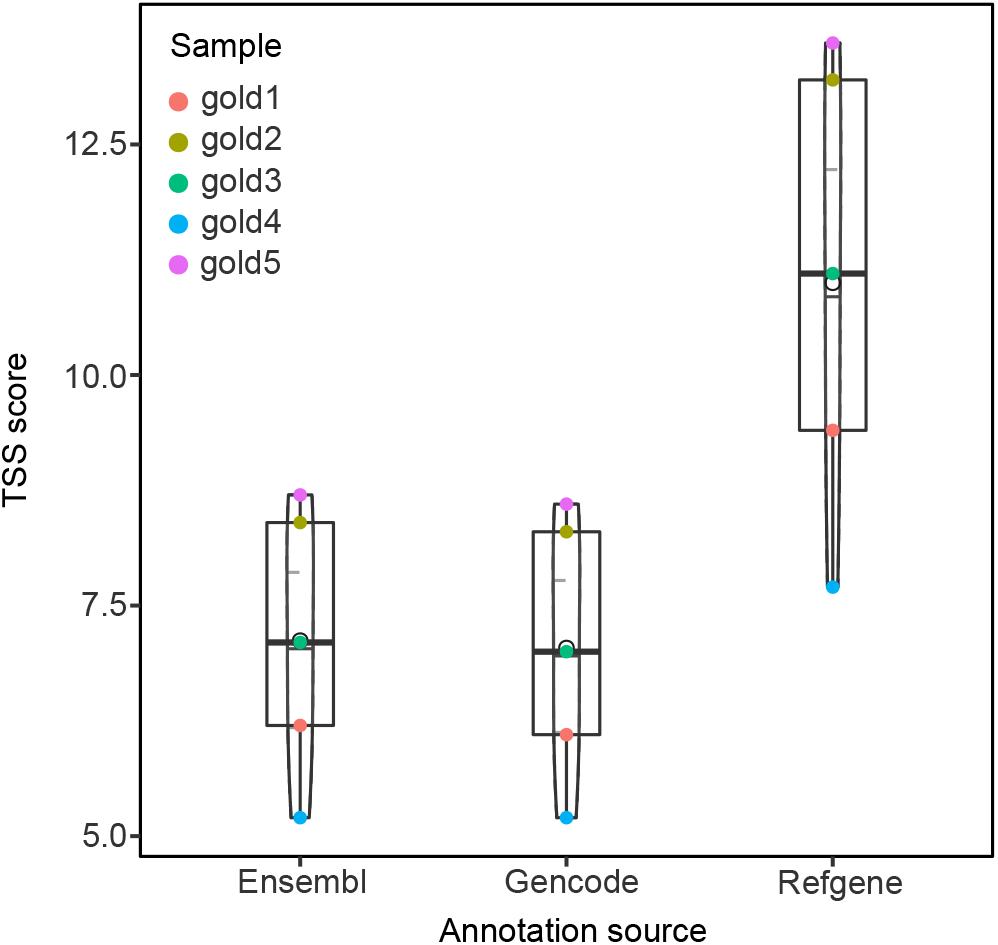
The TSS enrichment score is dependent on the annotation source. Refgene TSS annotations, which include the predominant TSS annotation only, produces the highest TSS enrichment score.

**Fig. S8:**
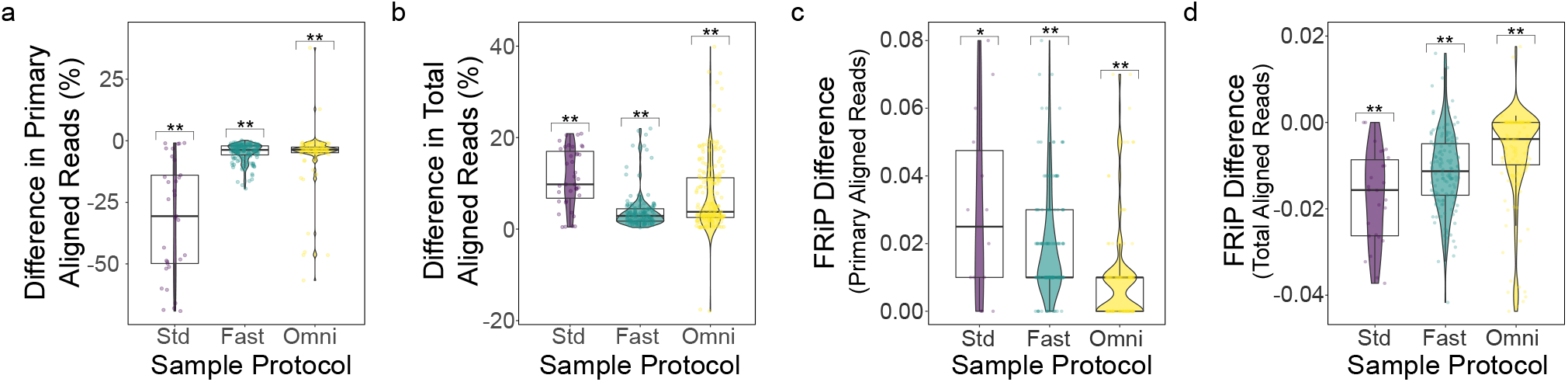
Prealignment changes the relationship between primary genome and total aligned reads and the fraction of reads in peaks (FRiP) is dependent on mapping strategy. (a) The number of primary, nuclear genome mapped reads is reduced when using prealignments. (b) However, the total number of mapped reads is increased with prealignments due to more specific read mapping. (c) The FRiP is increased with prealignments when using primary, nuclear genome mapped reads as the denominator. (d) In contrast, when using the total mapped reads the FRiP is reduced when using prealignments due to a larger mapped read pool in the denominator (* = p < 0.01; ** =p < 0.001; t-test (mu = 0) with Benjamini-Hochberg correction).

